# GCfix: A Fast and Accurate Fragment Length-Specific Method for Correcting GC Bias in Cell-Free DNA

**DOI:** 10.1101/2024.11.07.622399

**Authors:** Chowdhury Rafeed Rahman, Zhong Wee Poh, Anders Jacobsen Skanderup, Limsoon Wong

## Abstract

**Motivation:** Cell-free DNA (cfDNA) analysis has wide-ranging clinical applications due to its non-invasive nature. However, cfDNA fragmentomics and copy number analysis can be complicated by GC bias. There is a lack of GC correction software based on rigorous cfDNA GC bias analysis. Furthermore, there is no standardized metric for comparing GC bias correction methods across large sample sets, nor a rigorous experiment setup to demonstrate their effectiveness on cfDNA data at various coverage levels.

**Results:** We present GCfix, a method for robust GC bias correction in cfDNA data across diverse coverages. Developed following an in-depth analysis of cfDNA GC bias at the region and fragment length levels, GCfix is both fast and accurate. It works on all reference genomes and generates correction factors, tagged BAM files, and corrected coverage tracks. We also introduce two orthogonal performance metrics for (1) comparing the fragment count density distribution of GC content between expected and corrected samples, and (2) evaluating coverage profile improvement post-correction. GCfix outperforms existing cfDNA GC bias correction methods on these metrics.

**Availability:** GCfix software and code for reproducing the figures are publicly accessible on GitHub: https://github.com/Rafeed-bot/GCfix_Software.

## 1. Introduction

Cell-free DNA (cfDNA) is increasingly used as a biomarker in patient monitoring, disease detection, non-invasive prenatal testing, and transplant rejection assessment [12, 14, 17]. Its non-invasive collection from blood and the low cost of shallow whole-genome sequencing (WGS) make cfDNA appealing for clinical applications [15]. For instance, copy number alteration analysis and fragmentomics based on shallow WGS cfDNA are gaining traction in cancer diagnosis [1, 7, 30, 18] and tissue-of-origin detection [8, 19]. These analyses are also increasingly applied to deep WGS samples to quantify circulating tumor DNA (ctDNA) [29]. Such methods rely on counting reads or fragments across genomic regions. However, in practice, read counts from a given region depend on its GC content, with sequencing rates typically non-linearly reduced in GC- and AT-rich regions.

This GC bias in cfDNA data arises from biases in priming [11], size selection [21], PCR [16], and sequencing error likelihood [20], with PCR as the main contributor [2]. PCR-free library preparation is impractical for cfDNA due to low DNA input. Factors like sequencing technology, collection tube, processing timing, and library preparation kit further influence GC bias [5]. This variability hinders the generalizability of algorithms and machine learning models for cfDNA, underscoring the need for effective corrective measures. GC bias correction methods fall into two main categories: LOESS/bin-based and single-position methods. In LOESS methods, the genome is divided into bins, with fragment counts and GC contents calculated per bin. Equal-sized GC bins [26] or a smooth distribution of GC content [4] are assumed, and each genomic bin is assigned to an appropriate GC bin or GC range. The mean fragment count per GC bin or GC content is calculated, and a LOESS (Locally Estimated Scatterplot Smoothing) curve is generated based on GC bin or GC content vs mean coverage, representing the GC bias across different GC contents. Popular cfDNA tools, such as ichorCNA [1] (ctDNA quantification), Delfi [7] (cancer detection and tissue-of-origin identification) and WisecondorX [23] (non-invasive prenatal testing) use this approach. Ideally, bin sizes should be under 1 Kb, which is feasible only at high sequencing depths. For shallow WGS, larger bins are needed (due to fewer reads), which reduces accuracy; also, the GC bias curve is no longer bell-shaped and bins over 10 Kb rarely exceed 55% GC content [3, 9].

To address these limitations, a single-position GC bias correction method was introduced in [3]. This method uses a sliding GC context window across the reference genome to calculate expected fragment counts for different GC contents. A similar window is applied to each read in a sample, and observed counts are divided by the expected counts to obtain fragment rates for different GC contents. Initially developed for genomic DNA with a narrow fragment length range, this method uses a single GC context window for all fragment lengths. For cfDNA, which has varied fragment lengths, fragment length-specific GC context windows can better estimate GC bias. Griffin [8] applies this approach for cfDNA-based cancer subtyping but requires substantial runtime due to precise GC bias estimation across all positions in the reference genome and sample reads, and it lacks a standalone software package. GCparagon [25] is an approximation-based single-position method, trading some accuracy for significantly improved speed, and is available as a software. Both Griffin and GCparagon build on genomic DNA GC bias studies [3] but target cfDNA. Neither provides performance metrics for large sample evaluations, and both are limited to a single reference genome without comprehensive testing on cfDNA samples of varying coverage.

We propose GCfix, a cfDNA GC bias correction software based on precise, fragment length-specific single-position GC bias estimation. GCfix offers greater accuracy than Griffin, faster performance than GCparagon, and compatibility with all reference genomes. We introduce two orthogonal performance metrics for large-scale evaluation: the divergence metric, assessing corrected GC content fragment counts against expected distributions; and the variation metric, measuring coverage consistency across regions with similar copy numbers post-correction. GCfix is evaluated on four cohorts—two healthy [30], one colon, and one breast cancer [29]—across coverages from over 30X to 0.1X WGS. Finally, we show GCfix’s efficacy in enhancing ctDNA fraction prediction, fetal fraction estimation, and coverage consistency in gene promoter transcription start site (TSS) regions.

## 2. Materials and Methods

An overview of GCfix is provided in Fig. 1. Each step has been optimized for runtime and tailored based on the cfDNA GC bias analysis shown in Fig. 2 and Fig. S1-S3.

**Fig. 1.**
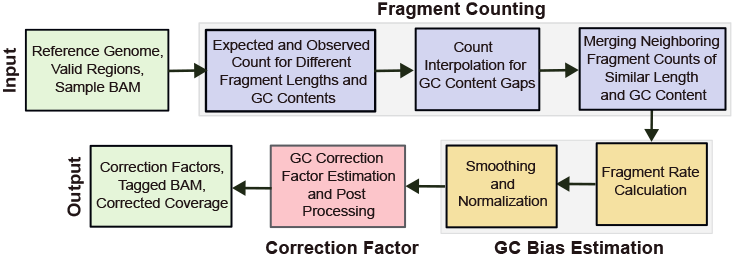
Overview of GCfix correction process.

**Fig. 2.**
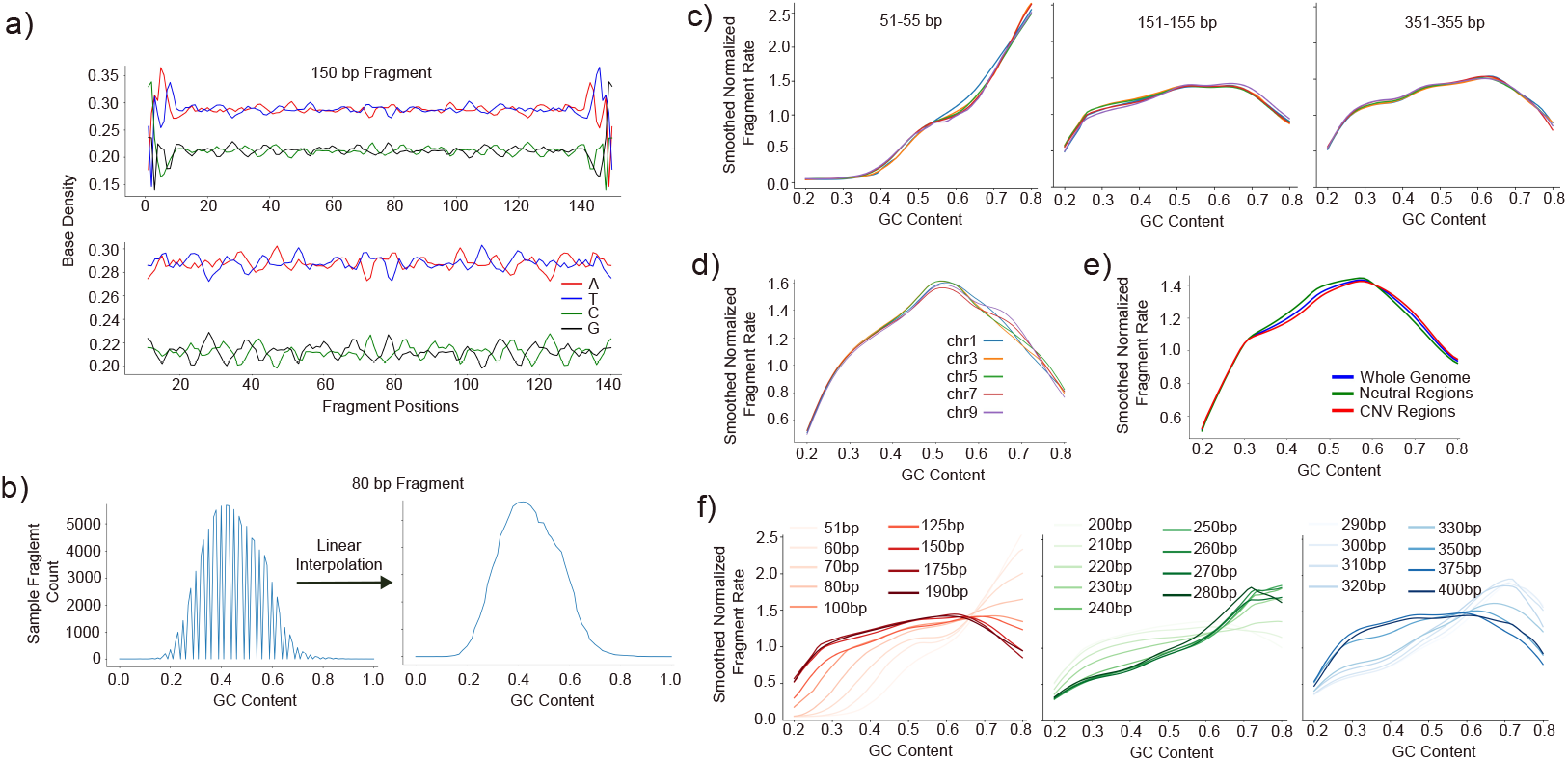
cfDNA GC bias investigation. (a) The noisy base density at start and end positions of 150 bp length fragments (top). The noise disappears when 10 bp is clipped from the start and end of the fragment (bottom). (b) Working with 101 fixed GC content bins irrespective of fragment length can render missing GC content bins for short fragments; this can easily be solved via linear interpolation. (c) Fragments within *±*2 bp length range generally exhibit similar GC bias curves for the same sample. (d) cfDNA GC bias in the same sample does not vary from chromosome to chromosome. (e) Estimated GC bias curves using whole genome, neutral regions and copy number variation regions look similar for the same cancer sample (f) There is a clear shift in fragment length GC bias curves while moving from short to long fragments of the same sample.

### 2.1. Input

GCfix requires a reference genome, valid genomic regions, and a sample’s BAM file as input. The reference genome provides expected fragment counts for different GC contents. Analysis is limited to valid genomic regions, excluding blacklisted, low mappability, low base-quality, and other marked unusual regions. The sample BAM file provides observed fragment counts across different GC contents by referencing sample reads mapped to the reference genome. By default, GCfix operates on fragments between 51-400 bp, which covers a majority of cfDNA fragments; other ranges can be specified as needed.

### 2.2. Fragment Counting

#### 2.2.1. Expected and Observed Fragment Count

We obtain counts for 101 GC content bins (0%, 1%, 2%, …, 100%) for fragments of length 51 to 400 bp (by default), resulting in a matrix of size 350 *×* 101 for both expected and observed settings. Fig. 2(a) shows noisy base density at the first and last 10 bp for fragments of length 150, a pattern consistently seen across all fragment lengths (Fig. S1) in both healthy and cancer samples. Thus, analogous to [3], for each fragment length, the GC context window is defined to exclude the first and last 10 bp of the fragment to avoid the noise in these positions. Only forward reads are counted, excluding duplicates, supplementary and unpaired reads. The GC content of each fragment is defined by dividing the count of G and C nucleotides in its GC context window by the window’s size. Fragments are then assigned into the GC content bins according to their length and GC content. For observed counts, the GC context windows of valid reads in the sample are used. For expected counts, the GC context window is slid across all valid positions in the reference genome (Suppl. Data 4).

#### 2.2.2. Count Interpolation for GC Content Gaps

As noted, we use 101 fixed GC content bins regardless of fragment length. This setup allows the expected and observed counts to be represented as uniform two-dimensional NumPy arrays, enabling parallel processing and SIMD (Single Instruction, Multiple Data) operations during GC bias estimation. However, using a fixed number of GC content bins can lead to gaps in observed and expected counts, especially for shorter fragments at smaller GC context sizes. The left plot of Fig. 2(b) displays observed fragment counts for 80 bp fragments at a 60 bp GC context size, where only 60 out of 101 GC content bins are filled. To address this, linear interpolation using neighboring non-zero GC content counts is applied, as shown in the right part of Fig. 2(b).

#### 2.2.3. Merging Neighboring Fragment Counts

When analyzing raw GC bias curves for different fragment lengths, we observe that fragments within *±*2 bp in length exhibit similar GC bias curves; cf. Fig. 2(c). For each fragment length, we merge its counts with those within *±*2 bp in the same GC content bin, both for observed and expected counts. This allows us to estimate accurate GC correction factors per fragment length, even in ultra-low-pass (ulp) WGS samples.

### 2.3. GC Bias Estimation

#### 2.3.1. Fragment Rate Calculation

Fragment rates are estimated for each GC content bin and fragment length by dividing the observed count by the expected count. We extract GC bias values for the 10%-90% GC content range, encompassing over 99.5% of the sample fragments. To minimize noise, we replace values below the 3^*rd*^ percentile with the 3^*rd*^ percentile, and values above the 97^*th*^ percentile with the 97^*th*^ percentile, of the obtained GC bias values.

#### 2.3.2. Normalization and Smoothing

We perform mean normalization on the denoised fragment rates across all GC content bins for each fragment length separately. Next, we apply LOESS smoothing using a smoothing factor of 0.1 for each fragment length and retain only the 20%-80% GC content range, which covers about 99% of the sample fragments. Previously, we extracted the 10% - 90% GC content range to assist the smoothing process for our target range, which is 20%-80%. Finally, we conduct mean normalization at the fragment length level one last time to obtain the final GC bias values.

Our analysis shows no major differences in chromosome-specific GC bias curves in the same sample, cf. Fig. 2(d), so we do not differentiate between genomic regions of the same sample during GC bias estimation. Although it was suggested [3] that including copy number variation (CNV) regions could confound GC bias estimation, we observe minimal differences in GC bias curves estimated across all fragments in CNV regions, neutral regions, and the whole genome for the same cancer sample; see Fig. 2(e). Similar analysis per fragment length confirms consistent GC bias curves for CNV, neutral, and whole genome regions (Fig. S2). This consistency across colon and breast cancer samples from two cohorts supports the inclusion of all valid regions, including CNV regions, in GC bias estimation.

Examining the GC bias curves for individual fragment lengths ranging from 51 to 400 bp, we observe a shift from a gradually increasing curve to bell curve after 80 bp, a transition from a bell curve to a gradually increasing curve after 220 bp, and another shift from a gradually increasing curve to a bell curve after 300 bp; cf. Fig. 2(f). This pattern is consistent across both healthy and cancer samples from different cohorts (Fig. S3), highlighting the importance of estimating GC bias at fragment length level for cfDNA data.

### 2.4. Correction Factor Estimation

Fragments with a GC bias value greater than 1 are overrepresented and should be down-weighted, while those with a GC bias value less than 1 should be up-weighted. Thus, the correction factor is the inverse of the estimated GC bias values for different fragment lengths and GC content bins. We estimate GC bias only for the 20% - 80% GC content range, setting correction factors for bins outside of this range to 0. To ensure that no correction factor exceeds 20, we apply a maximum threshold. However, if the 99^*th*^ percentile of correction factors for a sample exceeds 20, we set the maximum threshold to this percentile, assuming the sample has significant initial GC bias.

### 2.5. Output

GCfix generates three output files for a single input sample BAM file: (1) a CSV file containing GC bias correction factors for various fragment lengths (rows) and different GC contents (columns); (2) a tagged BAM file with the GC bias correction factor for each read as a ‘GC’ tag; and (3) a CSV file listing genomic bins and their corresponding corrected read counts.

## 3. Experiment Results

### 3.1. Dataset Description

All samples in this study are obtained from [30] and [29]. The evaluation plots in Fig. 3 and 4(b)-(c) represent four deep WGS (over 30X) cohorts: (a) two healthy cohorts with 12 and 17 samples; (b) one colon cancer cohort with 12 samples; and (c) one breast cancer cohort with 10 samples. These samples are downsampled to lower coverages (down to 0.1X) using SAMtools for GC bias correction testing in shallow WGS. Fig. 4(a) features lpWGS samples from 5 cohorts: (a) three healthy cohorts with 12, 17, and 28 samples; (b) 86 colon cancer samples; and (c) 10 breast cancer samples with available variant allele frequency (VAF).

**Fig. 3.**
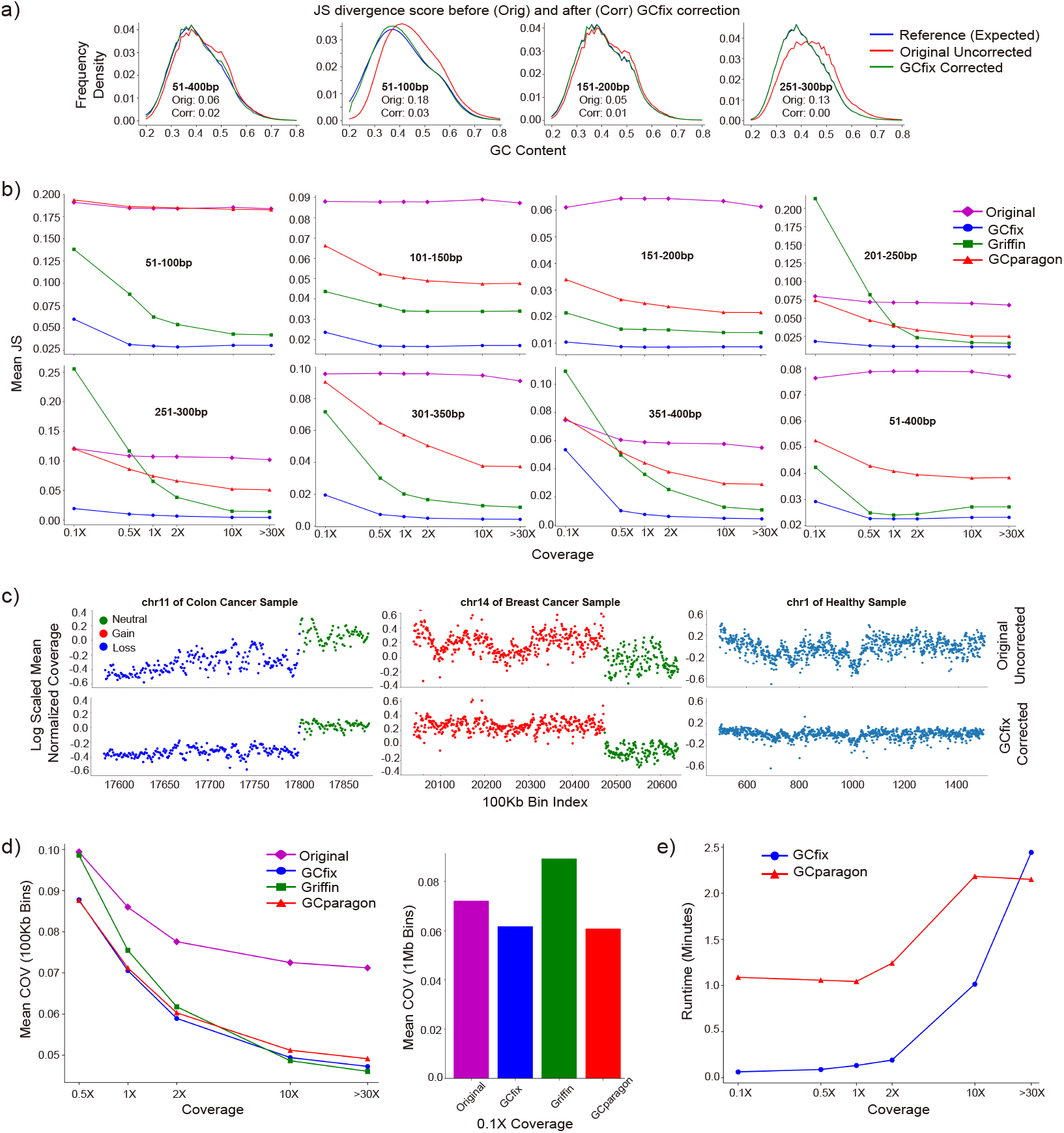
GCfix performance evaluation. (a) Shift in the GC content fragment frequency density distribution of original sample towards the expected distribution following GCfix correction for different fragment length groups. (b) Mean Jensen Shannon (JS) divergence from the expected GC content fragment frequency density distribution, across all samples, before (original) and after correction using three different methods across various coverages for different fragment lengths. (c) Coverage profile before and after GCfix correction for different contigs in cancer and healthy samples. (d) Mean coefficient of variation (COV) across similar CNV regions for all samples, before and after coverage profile correction using three different methods across various coverages for fragments ranging from 51 to 400 bp. The left and right subfigures use 100 Kb and 1 Mb bins in COV calculation, respectively. (e) Runtime comparison between GCfix and GCparagon for generating GC bias correction factors across samples of varying coverages.

**Fig. 4.**
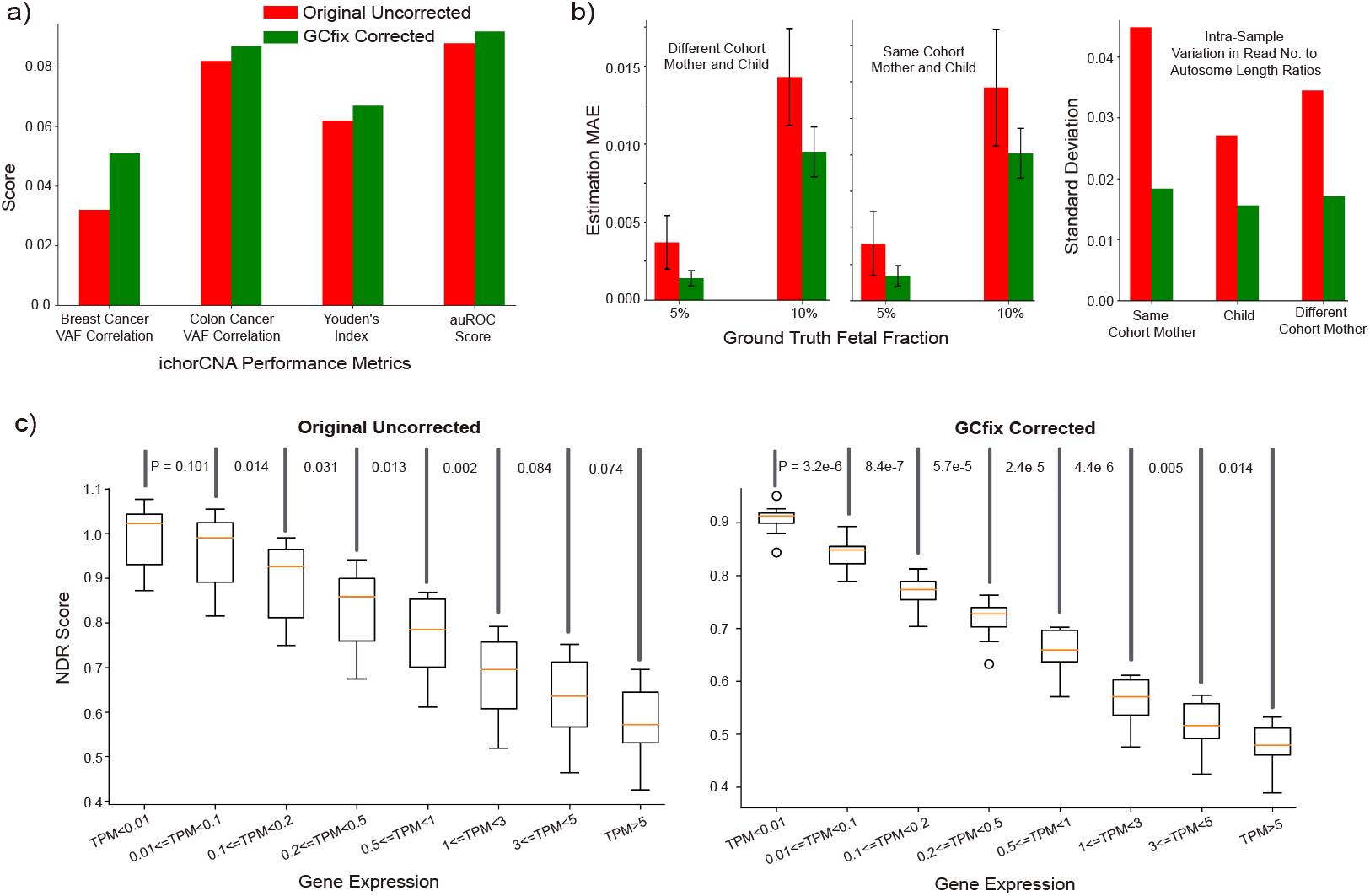
GCfix applications. (a) Enhanced ichorCNA performance in ctDNA quantification and cancer detection. (b) Improved simulated fetal fraction estimation. (c) Increased consistency of NDR score (derived from gene promoter TSS regions) across samples within the same gene expression group, and better separation between different gene expression groups.

### 3.2. Jensen Shannon (JS) Divergence

JS divergence measures ‘distance’ between two probability distributions. It is bounded between 0 and 1, where a lower score indicates greater similarity. The fragment frequency density across different GC contents for a sample represents a probability distribution. As shown in Fig. 3(a), the original uncorrected distribution deviates significantly from the expected reference distribution for various fragment length groups. In contrast, the corrected distribution aligns closely with the expected one, resulting in a very low JS divergence. Fig. 3(b) illustrates the mean JS divergence across all samples for different fragment length groups and coverages. GCfix significantly reduces divergence for all length groups and coverages compared to the original and other correction methods. Notably, the JS divergence for Griffin appears worse than uncorrected distributions for some length groups below 1X coverage. On the other hand, GCparagon shows minimal change in divergence scores for short and long fragments. Fig. S4 and S5 present boxplots comparing JS divergence scores for deep WGS and ulpWGS (0.1X) samples, confirming similar conclusions as in Fig. 3(b).

### 3.3. Coefficient of Variation (COV)

Bins within the same genomic region and sharing a similar copy number (CN) label (neutral/gain/loss) should exhibit comparable normalized coverage, barring any GC bias effects. Such a **run of bins** should have a low coefficient of variation (COV) for their normalized coverages, indicating that the standard deviation (std) of these coverages is low relative to their mean. For healthy samples, all bins are expected to be neutral regarding CNV. In cancer samples, we use only bins with high confidence CNV calls from ichorCNA. A CNV call is deemed high confidence under three conditions: (1) for neutral calls, CN should be between 1.5 and 2.5; (2) for gain/amplified/highly amplified calls, CN must exceed 2.5; (3) for homozygous/heterozygous deletion calls, CN should be below 1.5. We analyze runs of 100 Kb bins (1 Mb for 0.1X WGS) across the genome and calculate the COV for each run. The mean of these COVs is the sample COV. Each bin run has three properties: (a) it contains 50 consecutive 100 Kb bins (or 10 consecutive 1 Mb bins for 0.1X WGS); (b) all bins in a run share the same label (neutral/gain/loss); and (c) each run is at least 25 100 Kb bins (or 5 1 Mb bins for 0.1X WGS) apart from the immediate previous run.

Fig. 3(c) shows a significant reduction of COV for healthy and cancer samples in both neutral and CNV regions after GCfix correction. Fig. 3(d) displays the mean COV (across all samples) for original and corrected coverage profiles using different methods across various coverages. At 10X and higher coverage, Griffin and GCfix exhibit comparable mean COV, while GCparagon lags behind. However, Griffin performance drops significantly below 2X WGS, with its COV worse than the original for 0.1X WGS samples and similar to the original for 0.5X WGS. Fig. S13 visually shows how Griffin disrupts the original coverage profile for 0.1X WGS samples.

Low-pass WGS (lpWGS) cfDNA sequencing is widely used for analyzing genomic copy number alterations [1, 6, 13, 24], making effective coverage correction on lpWGS samples crucial. Fig. S6 and S7 are boxplots of COV scores for deep WGS and ulpWGS (0.1X) samples across the different correction methods, supporting the conclusions from Fig. 3(d). Fig. S6 shows COV boxplots for various fragment length groups, revealing that GCparagon does not improve COV for short fragments (51-100 bp). Fig. S12 provides visual evidence of this, comparing short fragment coverage profiles before and after GCparagon correction. GCfix performs well across all fragment lengths and coverages.

Runtime Comparison

GCfix is over 40 times faster than Griffin for both deep and lpWGS data (Fig. S9) while consuming a similar amount of memory (around 20 Mb). In contrast, GCparagon randomly samples around 5 million reads (equivalent to 0.5X WGS) for each sample, regardless of coverage, to estimate GC bias, which makes it extremely fast. Although GCfix uses all reads to accurately estimate GC bias, it is faster than GCparagon up to 30X deep WGS coverage; cf. Fig. 3(e).

## 4. GCfix Applications

### 4.1. ichorCNA Performance Improvement

cfDNA-based cancer detection [7, 18] and tumor fraction quantification [1, 30] have recently gained traction with lpWGS cfDNA data. This study examines the impact of GCfix on the ctDNA quantification software ichorCNA [1].

ichorCNA models copy number states across the genome using a Hidden Markov model (HMM) [22] on a wig file containing read count of 1 Mb bins. We generated a corrected wig file using GCfix for each sample and applied ichorCNA method to estimate the ctDNA fraction. As shown in Fig. 4(a), GCfix enhances the correlation between ichorCNA-estimated ctDNA fraction (based on lpWGS) and maximum VAF for both colon and breast cancer samples. Also, Youden’s index [27] and auROC score [10] for cancer and healthy sample discrimination are improved following GCfix correction (all relevant plots provided in Fig. S10).

### 4.2. Improvement in Fetal Fraction Estimation

Cost-effective non-invasive prenatal testing (NIPT) relies on lpWGS sequencing of maternal DNA [28]. We simulated fetal fraction estimation using two lpWGS BAM files—one for female and one for male. Each file was downsampled to have a similar number of fragments. We took X% reads from the male BAM file and merged them with the full female BAM file to create a new merged BAM representing a pregnant female where there was a male fetus inside with X% fetal fraction. The task here is to estimate this X% fetal fraction by analyzing read counts of each autosome inside the merged BAM file representing a pregnant female with a male fetus at X% fetal fraction.

We obtained 22 fetal fraction estimates (detailed in Suppl. Data 1) from the 22 autosome pairs and calculated their mean as the final estimate. We tested healthy female (mother) and male (fetus) samples from both the same cohort and different cohorts, varying X% as 5% and 10% to represent the ground truth fetal fraction. Fig. 4(b) (left) shows the improvement in mean absolute error (MAE) of estimated fetal fractions after applying GCfix correction to the female pregnant BAM. The std of the 22 estimations also significantly decreased post-correction. In Fig. 4(b) (right), we analyzed the mapped read count-to-autosome length ratio for all 22 autosomes within each sample, calculating std (ideally, std should be 0). The results indicate that after GCfix correction, the std reduces by around 50% for all samples, explaining the improved estimation accuracy.

### 4.3. Improvement in NDR Coverage Profile

The relative coverage at nucleosome-depleted regions (NDRs) around transcription start sites (TSSs) is associated with gene expression. We examine the coverage profile and NDR score [29] of gene promoter TSS regions obtained from the Matched Annotation from NCBI and EMBL-EBI (MANE) database. NDRs are defined as the −150 to +50 bp window around the TSS, while flanking regions span −1000 bp to −2000 bp upstream and 1000 bp to 2000 bp downstream. Each TSS’s NDR score is calculated by dividing the mean coverage within the TSS region by the mean coverage in the flanking regions.

We analyzed NDR scores for 17 deep WGS healthy cfDNA TSS regions across gene expression levels from GTEx Whole Blood RNA-Seq before and after GCfix correction; cf. Fig. 4(c). Each boxplot point represents the mean NDR score of all NDRs in a single sample for a given gene expression group measured in TPM (Transcripts Per Million). As shown, NDR score variance within each expression group decreases substantially post-GCfix, yielding a statistically significant lower NDR score for each higher expression group compared to the immediate previous expression group (one-sided Wilcoxon Rank Sum test). Further, examining relative coverage (Suppl. Data 2) of *±*2000 bp region relative to each TSS position, GCfix correction markedly reduces variance in relative coverage profile within each gene expression group for the deep WGS healthy cfDNA samples (Fig. S11). The potential for NDR coverage improvement through GC correction was first discussed by Wee et al. for deep WGS colon cancer samples (manuscript in preparation).

## 5. Discussion and Concluding Remarks

In [3], single-position GC bias correction methods outperformed LOESS/bin-based methods, leading to Griffin’s fragment-length-specific single-position method for cfDNA GC bias correction. However, Griffin has a high runtime (see Section 3.4), performs poorly on shallow sequencing data, and is not user-friendly. It requires users to generate precomputed GC frequency files for their reference genome and lacks a tagged BAM output for convenient correction factor access. Users must consult multiple files and the read GC context to retrieve correction factors. Also, Griffin often produces NAN and 0 values as the estimated GC bias for some GC count and fragment length combinations due to insufficient post-processing of GC bias estimates. GCparagon builds on Griffin’s principles but uses sampling-based approximation to speed up correction factor generation, and is available as a software specifically for hg19. While it provides BAM tagging with correction factors, the sequential tagging process is slow. GCparagon’s weakness is its limited correction for short and long cfDNA fragments; cf. the flat GCparagon curves in Fig. S14. Fragment counts before and after GCparagon correction change by less than 1% (Fig. S8) because its correction factors are near 1, indicating conservative correction. As GCparagon estimates GC bias from a small subset of reads regardless of coverage, its GC bias correction performance on moderately high-coverage samples is weaker than both GCfix and Griffin. Algorithmic differences between GCfix, Griffin, and GCparagon are detailed in Suppl. Data 3.

GCfix addresses the issues above by efficiently using all reads in a sample BAM file to accurately estimate GC bias. It outperforms both Griffin and GCparagon across different sequencing coverages, as demonstrated using two performance metrics proposed here—JS divergence and COV. GCfix is also effective in applications such as ctDNA quantification, fetal fraction estimation, and TSS coverage profile improvement. Designed for ease of use, GCfix is compatible with all reference genomes and tags BAM reads with correction factors quickly (about 5 minutes for 1X WGS) by breaking BAM files into chunks, and tagging them in parallel. Users can specify the genomic regions for tagging, enhancing efficiency, or request corrected read counts for specific regions without needing expertise in GC bias correction. We expect GCfix to have broad applications across various research and clinical domains.

## References

1. V. A. Adalsteinsson, G. Ha, S. S. Freeman, et al. Scalable whole-exome sequencing of cell-free dna reveals high concordance with metastatic tumors. Nature communications, 8(1):1324, 2017.

2. D. Aird, M. G. Ross, W.-S. Chen, et al. Analyzing and minimizing pcr amplification bias in illumina sequencing libraries. Genome biology, 12:1–14, 2011.

3. Y. Benjamini and T. P. Speed. Summarizing and correcting the gc content bias in high-throughput sequencing. Nucleic acids research, 40(10):e72–e72, 2012.

4. V. Boeva, A. Zinovyev, K. Bleakley, et al. Control-free calling of copy number alterations in deep-sequencing data using gc-content normalization. Bioinformatics, 27(2):268–269, 2011.

5. P. D. Browne, T. K. Nielsen, W. Kot, et al. Gc bias affects genomic and metagenomic reconstructions, underrepresenting gc-poor organisms. GigaScience, 9(2):giaa008, 2020.

6. X. Chen, C.-W. Chang, J. M. Spoerke, et al. Low-pass whole-genome sequencing of circulating cell-free dna demonstrates dynamic changes in genomic copy number in a squamous lung cancer clinical cohort. Clinical Cancer Research, 25(7):2254–2263, 2019.

7. S. Cristiano, A. Leal, J. Phallen, et al. Genome-wide cell-free dna fragmentation in patients with cancer. Nature, 570(7761):385–389, 2019.

8. A.-L. Doebley, M. Ko, H. Liao, et al. A framework for clinical cancer subtyping from nucleosome profiling of cell-free dna. Nature Communications, 13(1):7475, 2022.

9. J. C. Dohm, C. Lottaz, T. Borodina, and H. Himmelbauer. Substantial biases in ultra-short read data sets from high-throughput dna sequencing. Nucleic acids research, 36(16):e105, 2008.

10. J. A. Hanley and B. J. McNeil. The meaning and use of the area under a receiver operating characteristic (roc) curve. Radiology, 143(1):29–36, 1982.

11. K. D. Hansen, S. E. Brenner, and S. Dudoit. Biases in illumina transcriptome sequencing caused by random hexamer priming. Nucleic acids research, 38(12):e131– e131, 2010.

12. E. Heitzer, I. S. Haque, C. E. Roberts, and M. R. Speicher. Current and future perspectives of liquid biopsies in genomics-driven oncology. Nature Reviews Genetics, 20(2):71–88, 2019.

13. D. H. Hovelson, C.-J. Liu, Y. Wang, et al. Rapid, ultra low coverage copy number profiling of cell-free dna as a precision oncology screening strategy. Oncotarget, 8(52):89848, 2017.

14. M. Ignatiadis, G. W. Sledge, and S. S. Jeffrey. Liquid biopsy enters the clinical: implementation issues and future challenges. Nature reviews Clinical oncology, 18(5):297–312, 2021.

15. P. Jiang and Y. M. D. Lo. The long and short of circulating cell-free dna and the ins and outs of molecular diagnostics. Trends in Genetics, 32(6):360–371, 2016.

16. I. Kozarewa, Z. Ning, M. A. Quail, et al. Amplification-free illumina sequencing-library preparation facilitates improved mapping and assembly of (g+ c)-biased genomes. Nature methods, 6(4):291–295, 2009.

17. Y. M. D. Lo, D. S. C. Han, P. Jiang, and R. W. K. Chiu. Epigenetics, fragmentomics, and topology of cell-free dna in liquid biopsies. Science, 372(6538):eaaw3616, 2021.

18. F. Mouliere, D. Chandrananda, A. M. Piskorz, et al. Enhanced detection of circulating tumor dna by fragment size analysis. Science translational medicine, 10(466):eaat4921, 2018.

19. F. Mouliere, B. Robert, E. Arnau Peyrotte, et al. High fragmentation characterizes tumour-derived circulating dna. PloS one, 6(9):e23418, 2011.

20. K. Nakamura, T. Oshima, T. Morimoto, et al. Sequence-specific error profile of illumina sequencers. Nucleic acids research, 39(13):e90–e90, 2011.

21. M. A. Quail, I. Kozarewa, F. Smith, et al. A large genome center’s improvements to the illumina sequencing system. Nature methods, 5(12):1005–1010, 2008.

22. L. R. Rabiner. A tutorial on hidden markov models and selected applications in speech recognition. Proceedings of the IEEE, 77(2):257–286, 1989.

23. L. Raman, A. Dheedene, M. De Smet, et al. Wisecondorx: improved copy number detection for routine shallow whole-genome sequencing. Nucleic acids research, 47(4):1605–1614, 2019.

24. J. Smolander, S. Khan, K. Singaravelu, et al. Evaluation of tools for identifying large copy number variations from ultra-low-coverage whole-genome sequencing data. BMC genomics, 22(1):357, 2021.

25. B. Spiegl, F. Kapidzic, S. Röner, et al. Gcparagon: evaluating and correcting gc biases in cell-free dna at the fragment level. NAR Genomics and Bioinformatics, 5(4):lqad102, 2023.

26. S. Yoon, Z. Xuan, V. Makarov, et al. Sensitive and accurate detection of copy number variants using read depth of coverage. Genome research, 19(9):1586–1592, 2009.

27. W. J. Youden. Index for rating diagnostic tests. Cancer, 3(1):32–35, 1950.

28. D. Yu, K. Zhang, M. Han, et al. Noninvasive prenatal testing for fetal subchromosomal copy number variations and chromosomal aneuploidy by low-pass whole-genome sequencing. Molecular genetics & genomic medicine, 7(6):e674, 2019.

29. G. Zhu, Y. A. Guo, D. Ho, et al. Tissue-specific cell-free dna degradation quantifies circulating tumor dna burden. Nature communications, 12(1):2229, 2021.

30. G. Zhu, C. R. Rahman, V. Getty, et al. Fragle: Universal ctdna quantification using deep learning of fragmentomic profiles. bioRxiv, pages 2023–07, 2023.

