## Supplementary Figures for "GCfix: A Fast and Accurate Fragment Length-Specific Method for Correcting GC Bias in Cell-Free DNA"

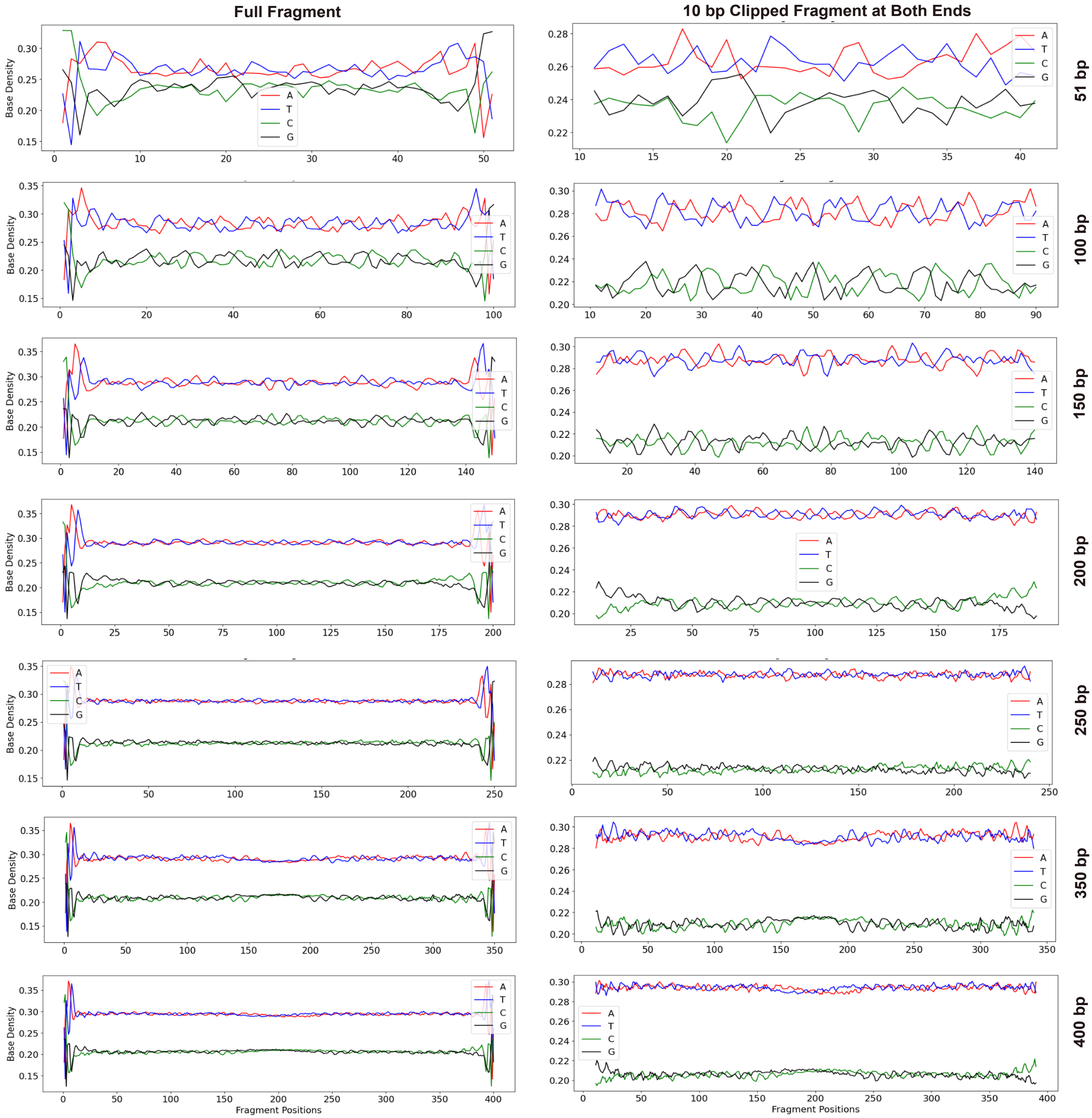


**Suppl Fig S1:** Base density in reference genome for full fragment GC context vs 10bp clipped (from both ends) fragment GC context for different fragment lengths of a healthy sample

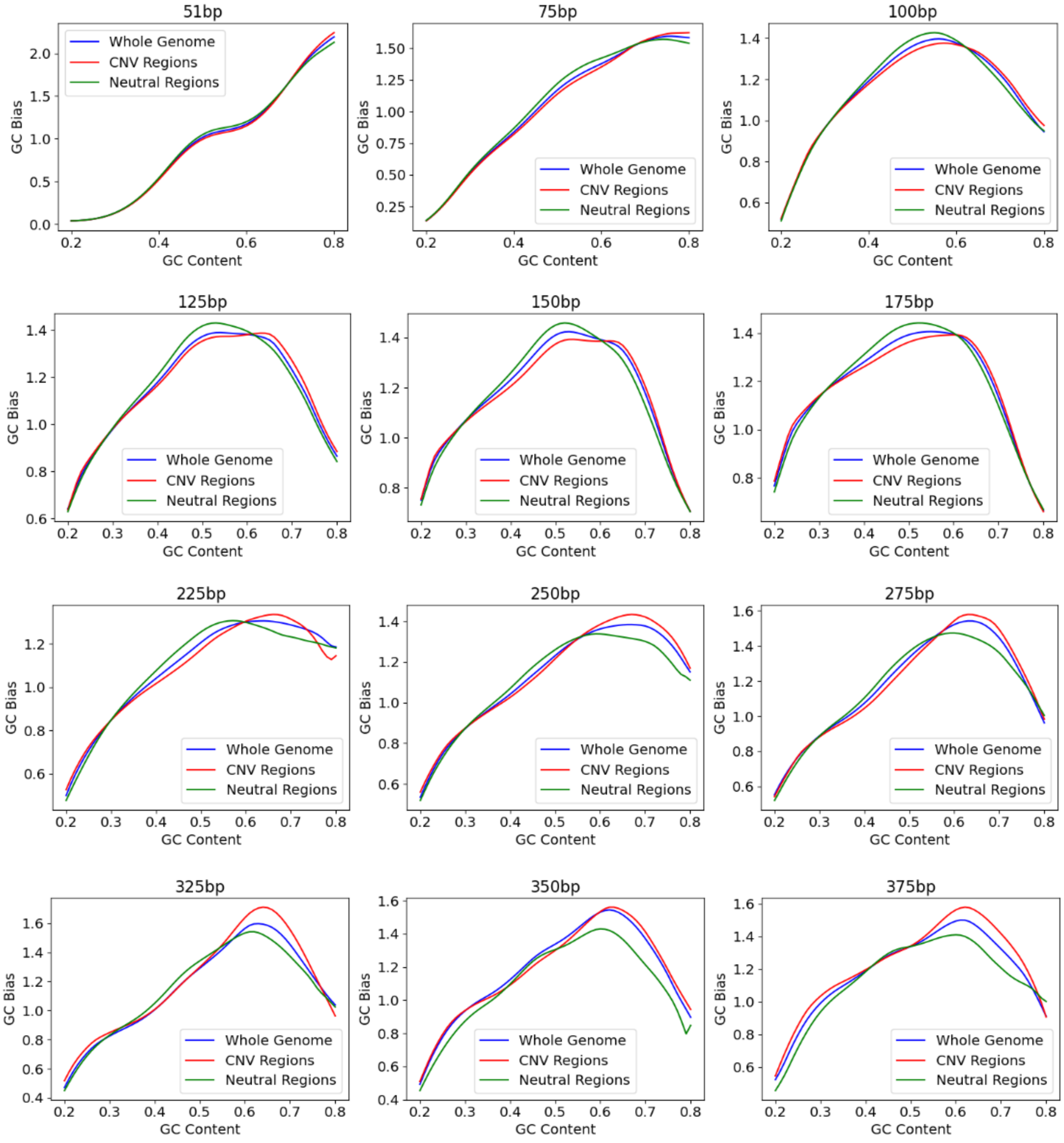


**Suppl Fig S2:** GC bias curve based on whole genome, neutral and copy number variant (CNV) regions for different fragment lengths of a colon cancer sample

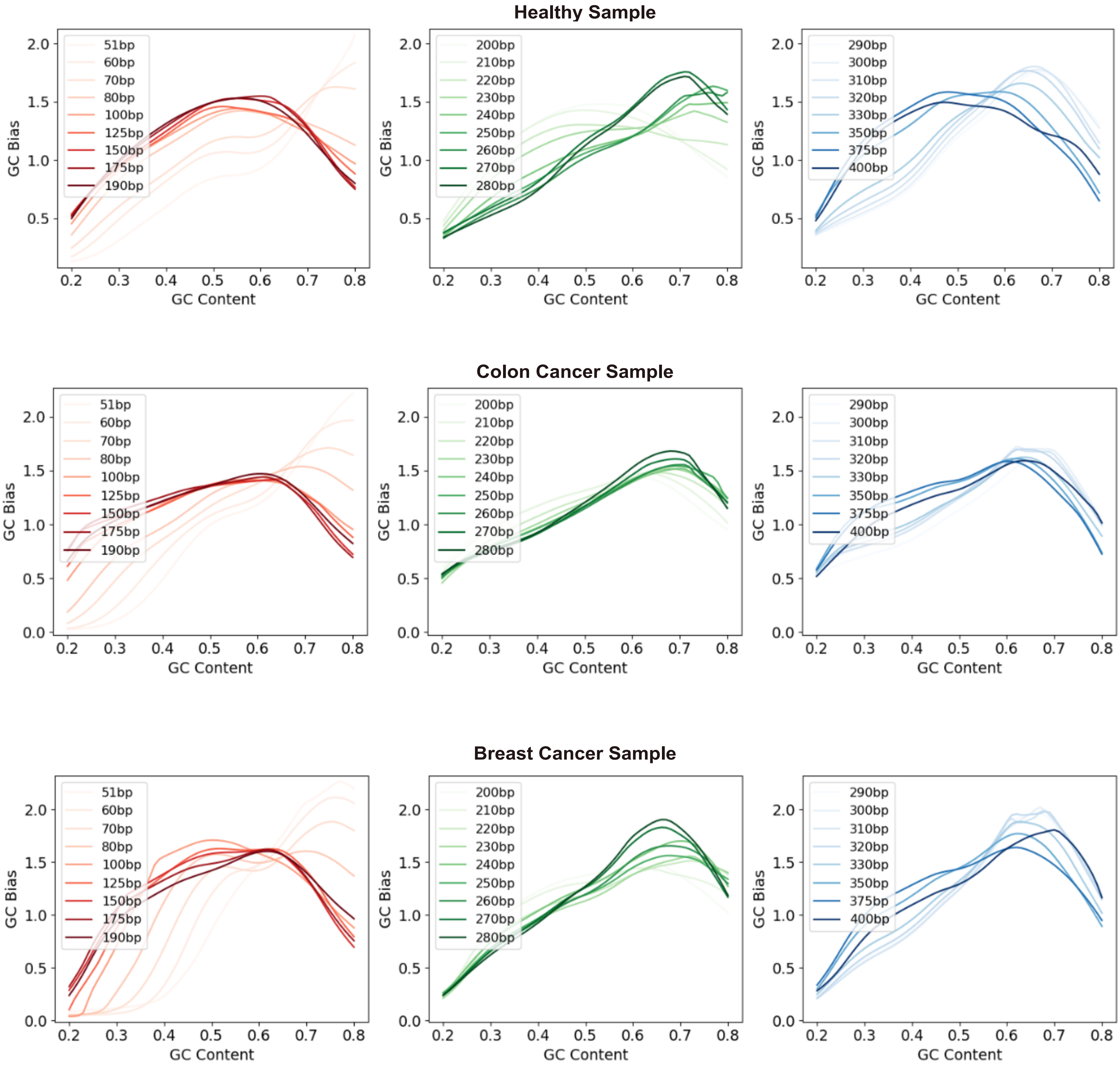


**Suppl Fig S3:** GC bias curves for different fragment lengths for healthy and cancer samples from different cohorts

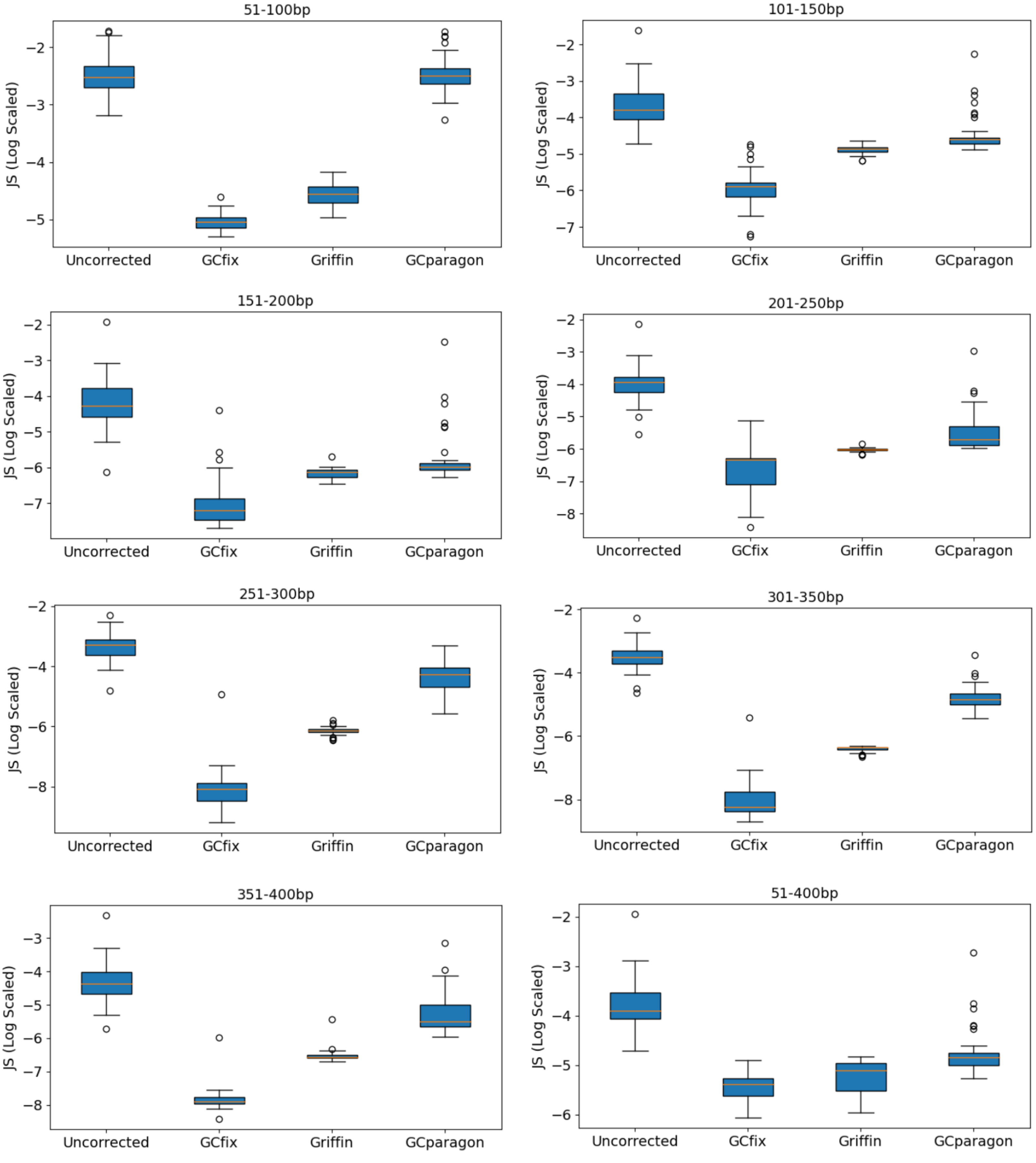


**Suppl Fig S4:** Sample JS divergence (log scaled) before and after correction (using different methods) from expected GC content fragment count density distribution based on different fragment length groups. 51 **deep WGS (>30X)** samples (both healthy and cancer) from 4 different cohorts have been used for each box plot.

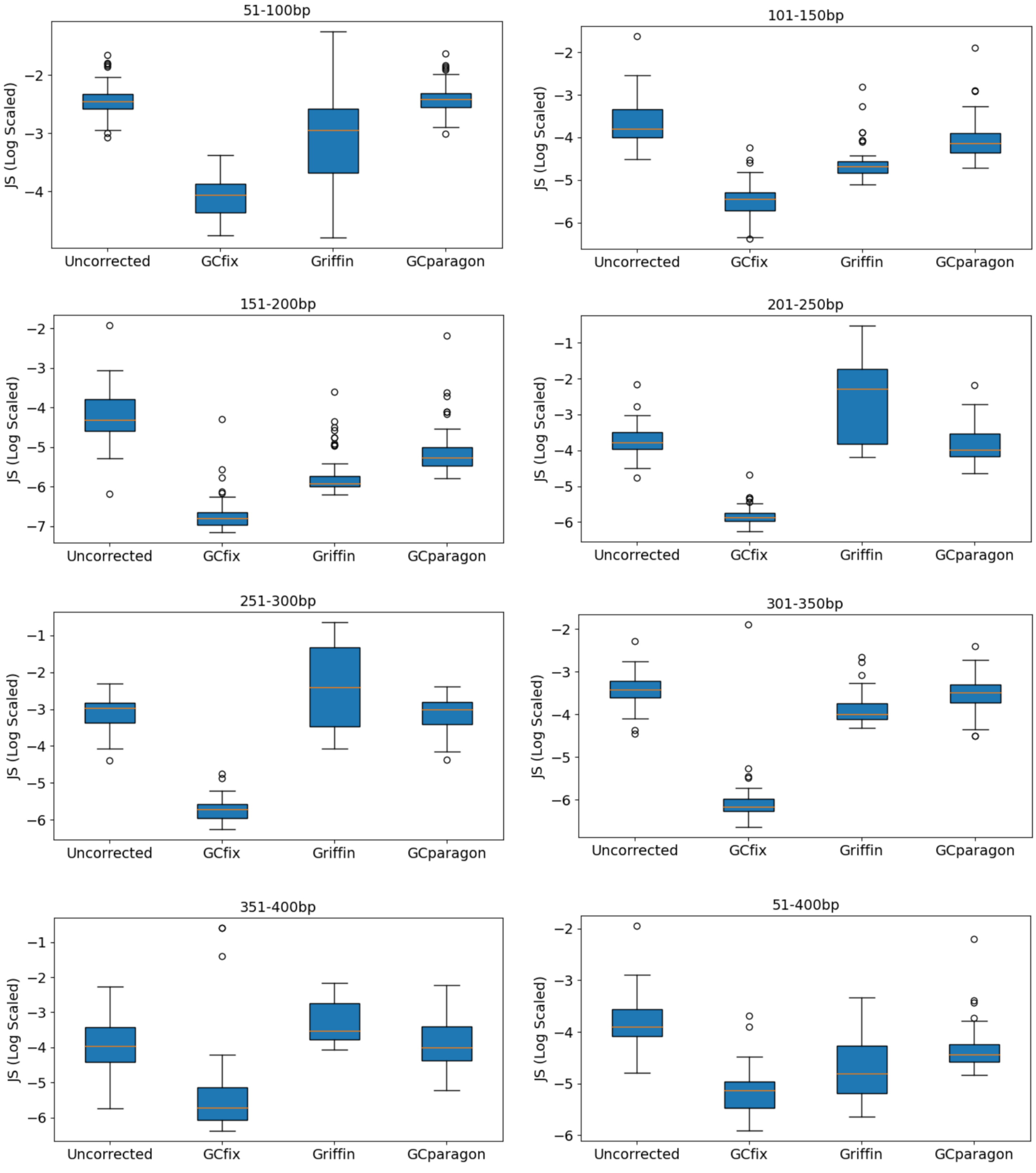


**Suppl Fig S5:** Sample JS divergence (log scaled) before and after correction (using different methods) from expected GC content fragment count density distribution based on different fragment length groups. 51 **ultra low pass WGS (~0.1X)** samples (both healthy and cancer) from 4 different cohorts have been used for each box plot.

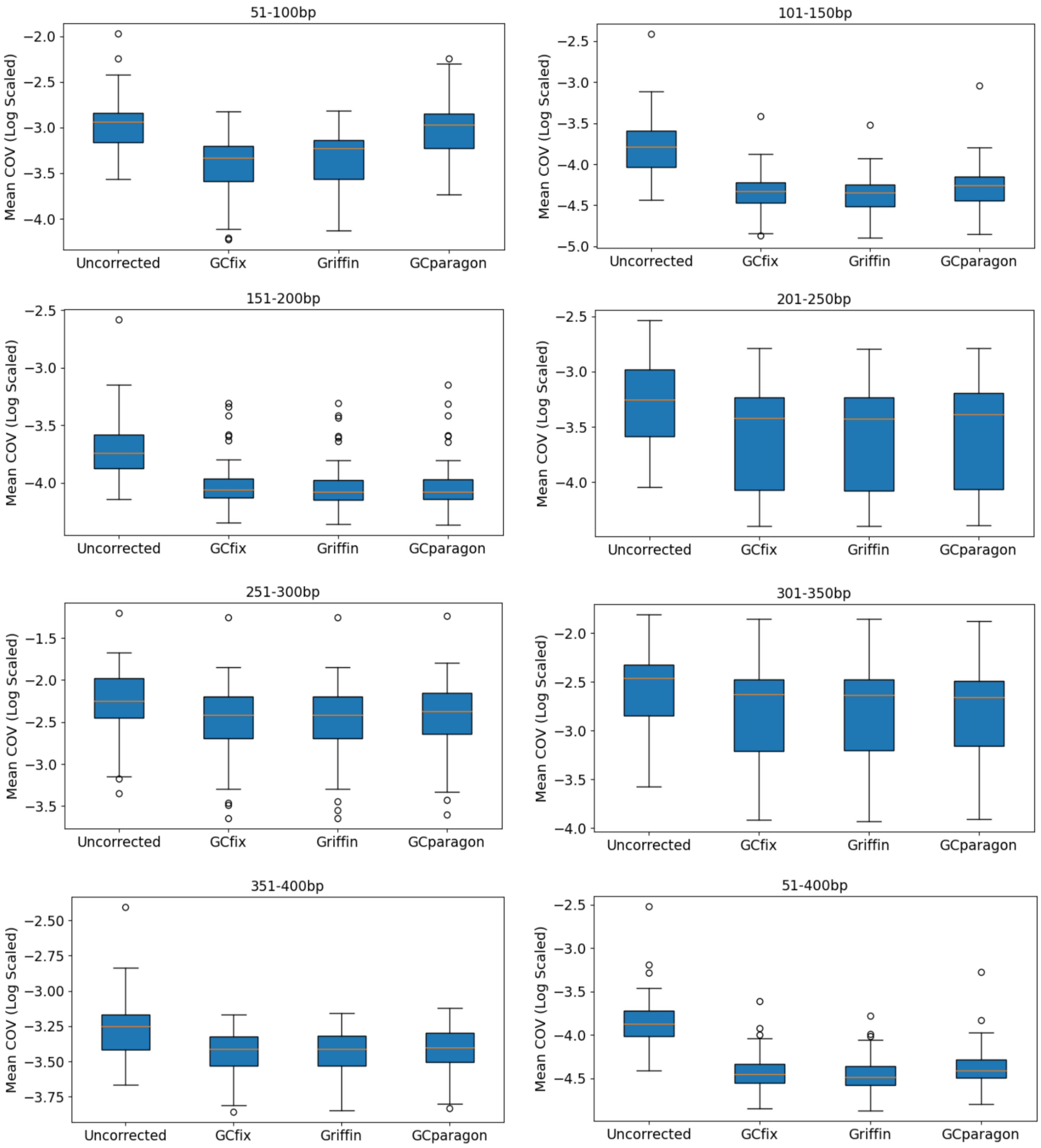


**Suppl Fig S6:** Sample mean coefficient of variation (log scaled) before and after correction (using different methods) based on different fragment length groups. 51 **deep WGS (>30X)** samples (both healthy and cancer) from 4 different cohorts have been used for each box plot.

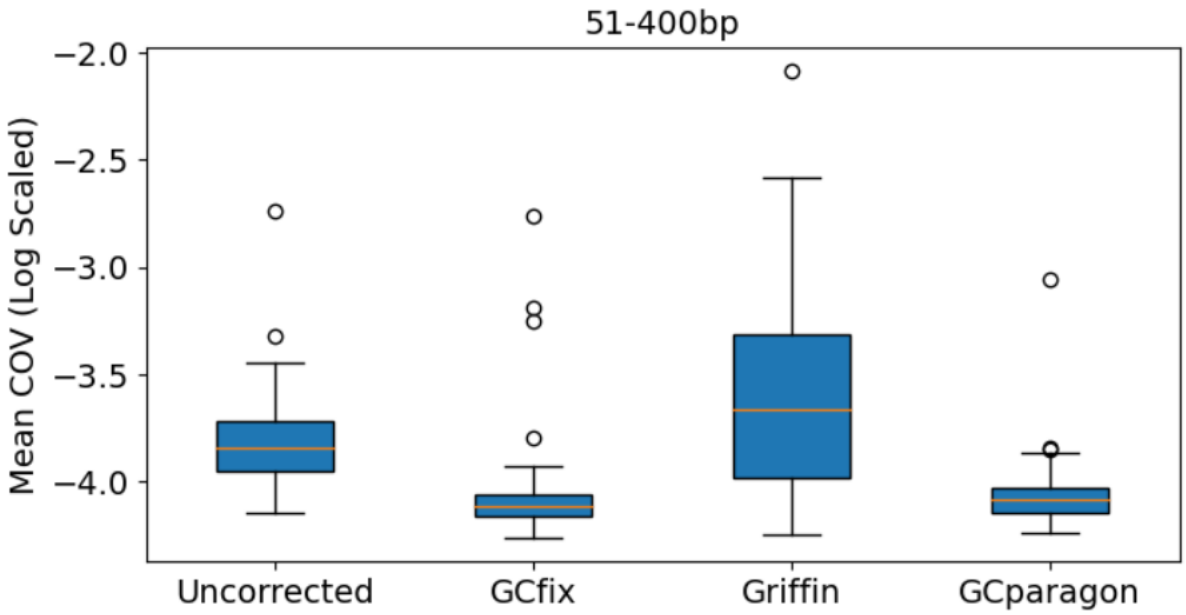


**Suppl Fig S7:** Sample mean coefficient of variation (log scaled) before and after correction (using different methods) based on all fragments of length 51-400bp. 51 **ultra low pass WGS (~0.1X)** samples (both healthy and cancer) from 4 different cohorts have been used for each box plot.

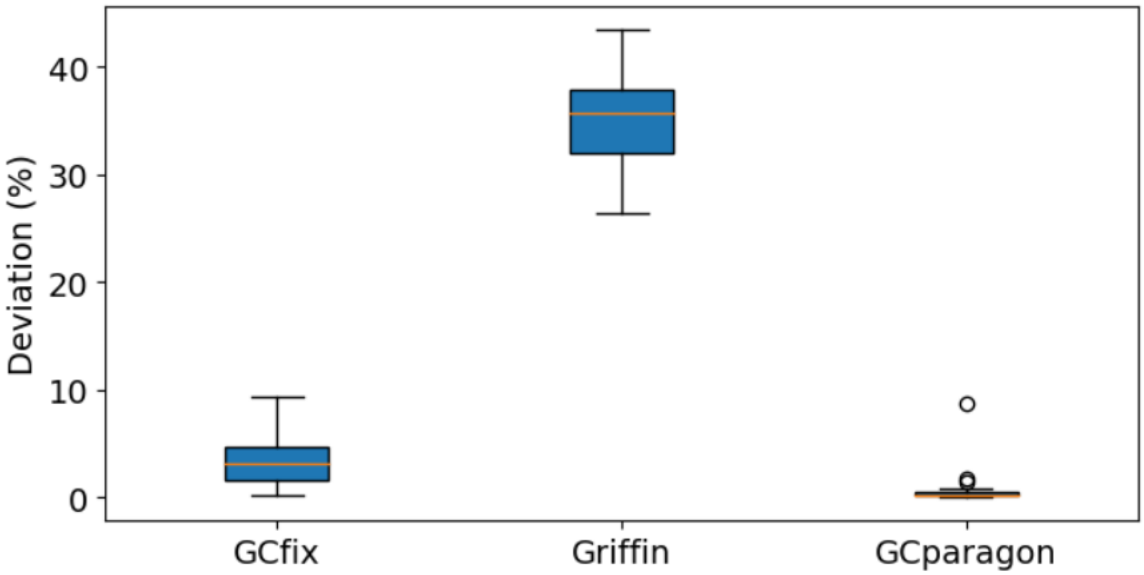


**Suppl Fig S8:** Deviation (%) in total fragment number after correction (using different methods) relative to the original uncorrected sample total fragment number using all samples (both healthy and cancer) from 4 different cohorts

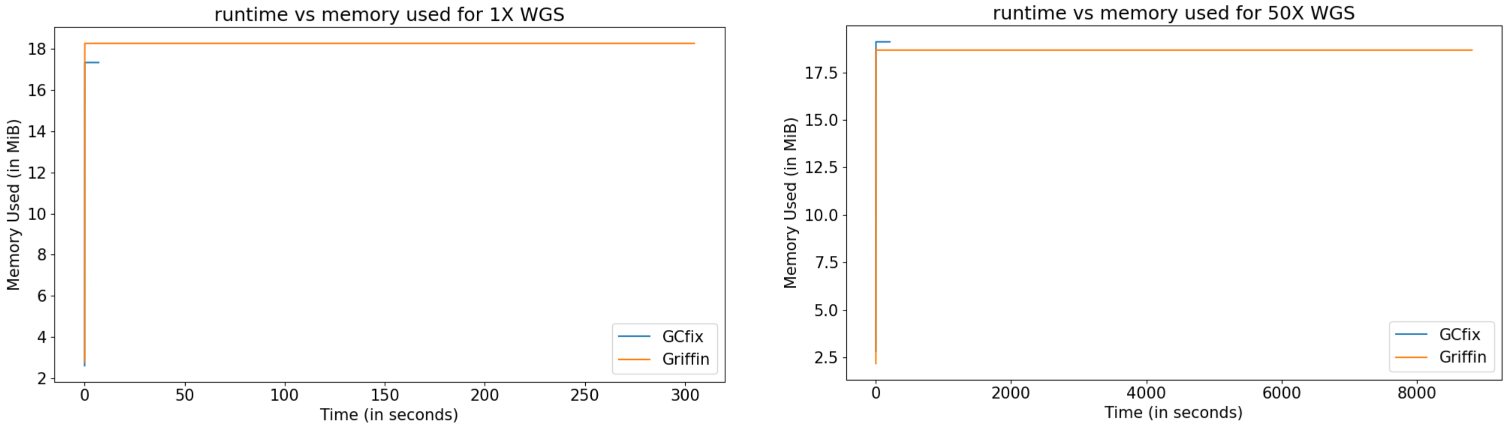


**Suppl Fig S9:** Runtime and memory comparison between GCfix and Griffin for 1X and 50X WGS sample

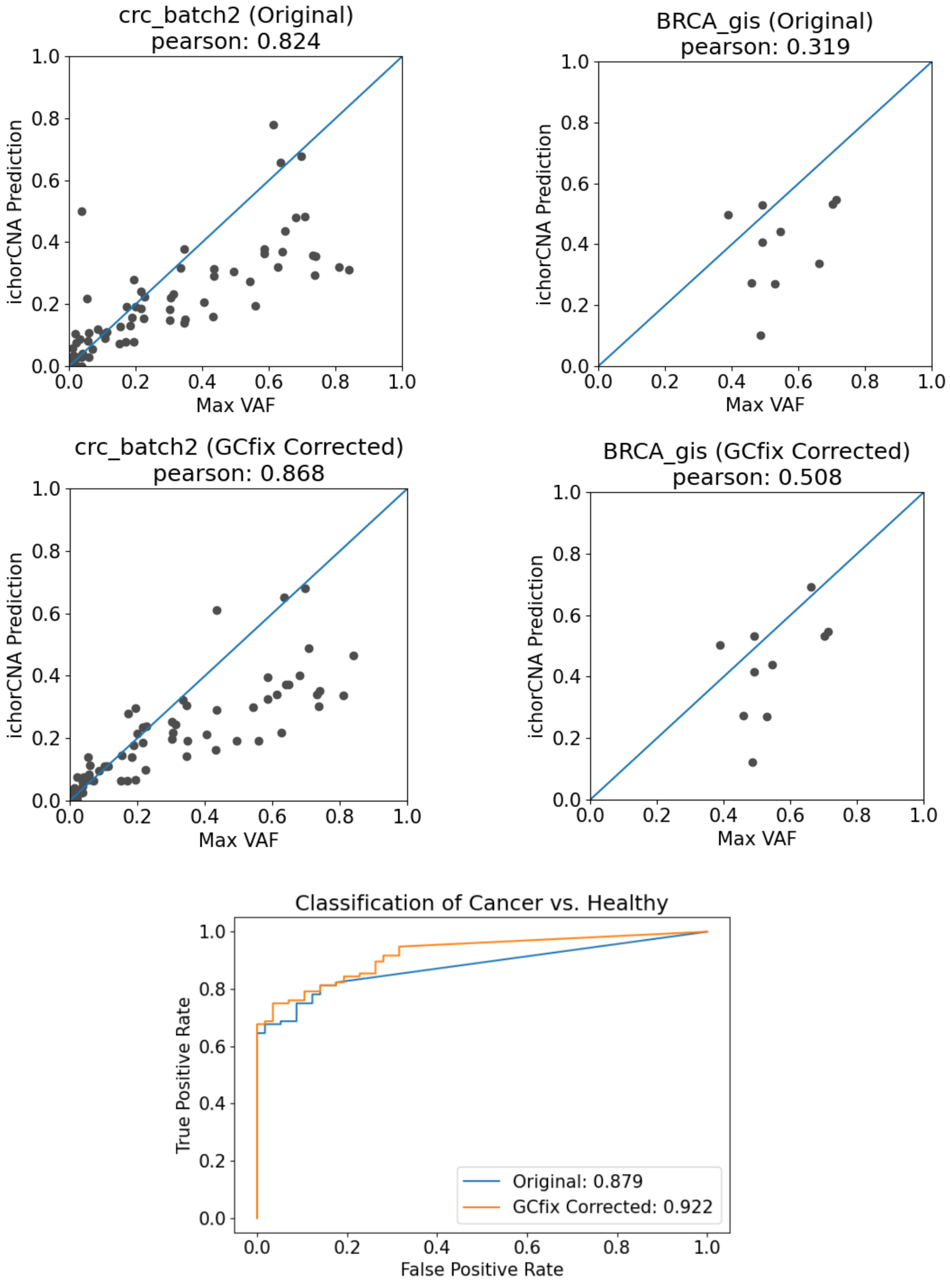


**Suppl Fig S10:** ichorCNA max VAF correlation and auROC plot before and after GCfix correction

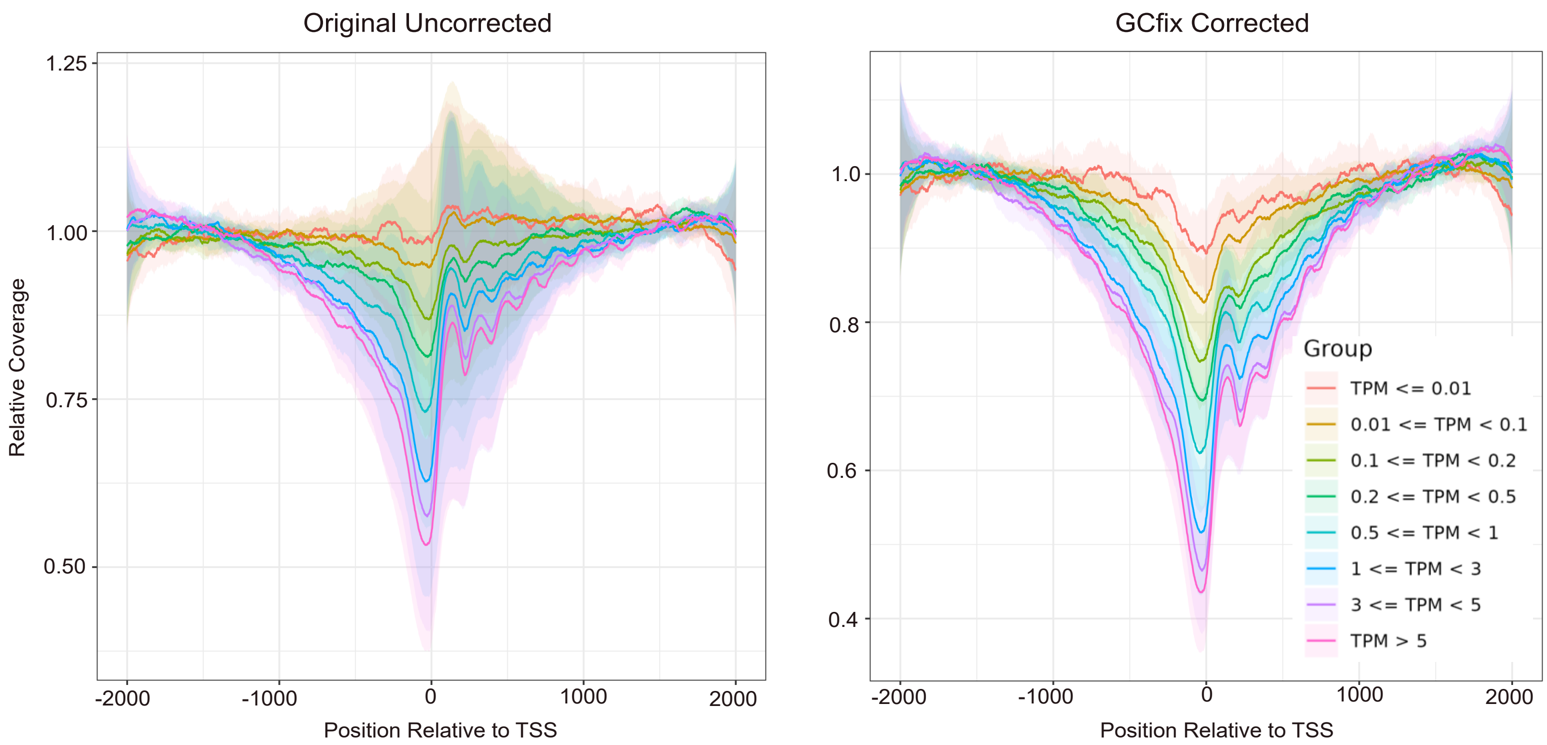


**Suppl Fig S11:** Relative coverage profile for different gene expression group nucleosome depleted regions (NDR) for the 17 deep WGS healthy samples before and after GCfix correction. GC correction significantly reduces the inter-sample variance in relative coverage profiles for the different gene expression groups.

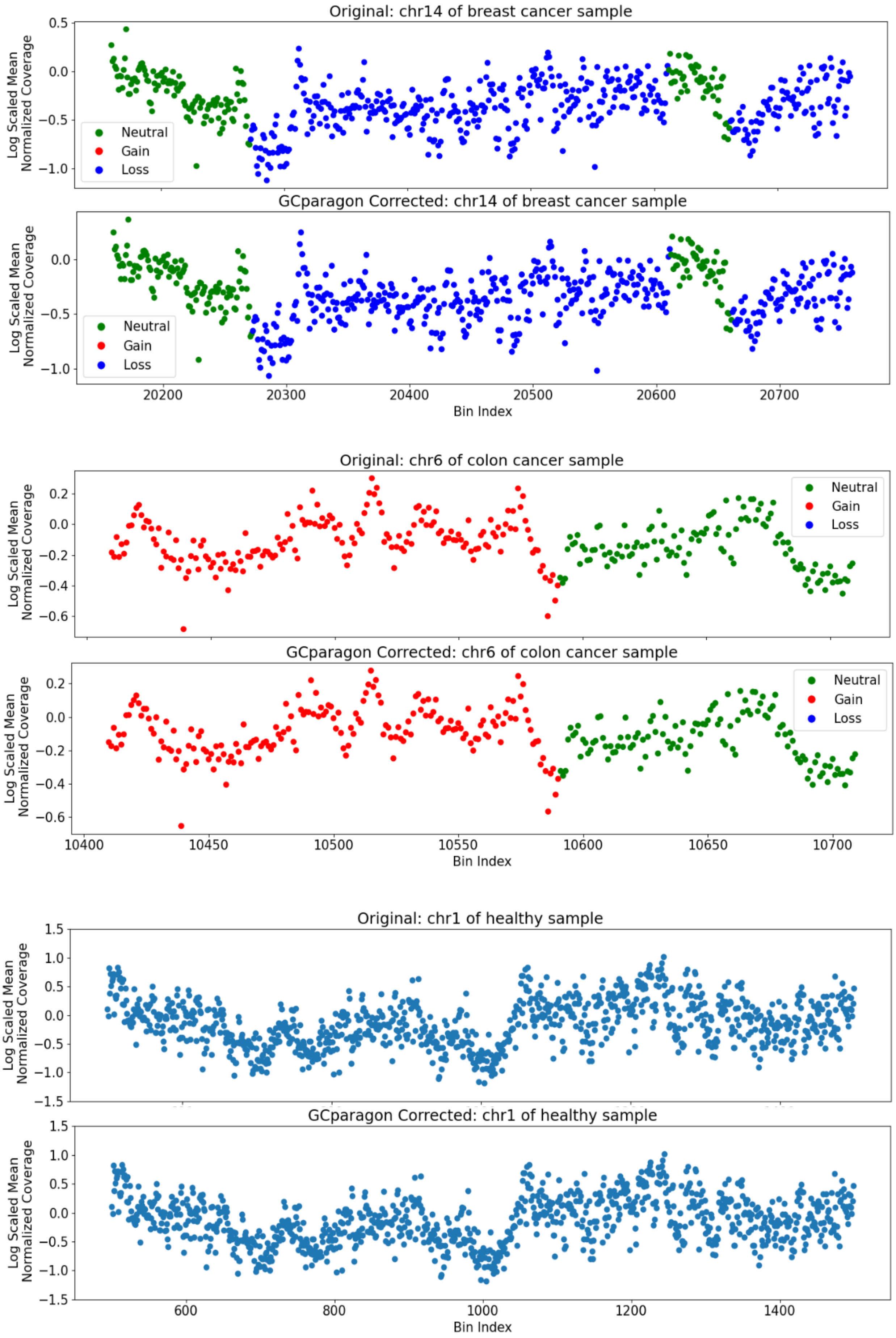


**Suppl Fig S12:** GCparagon does not correct for short fragment (51-100bp) coverage profile for cancer and healthy samples

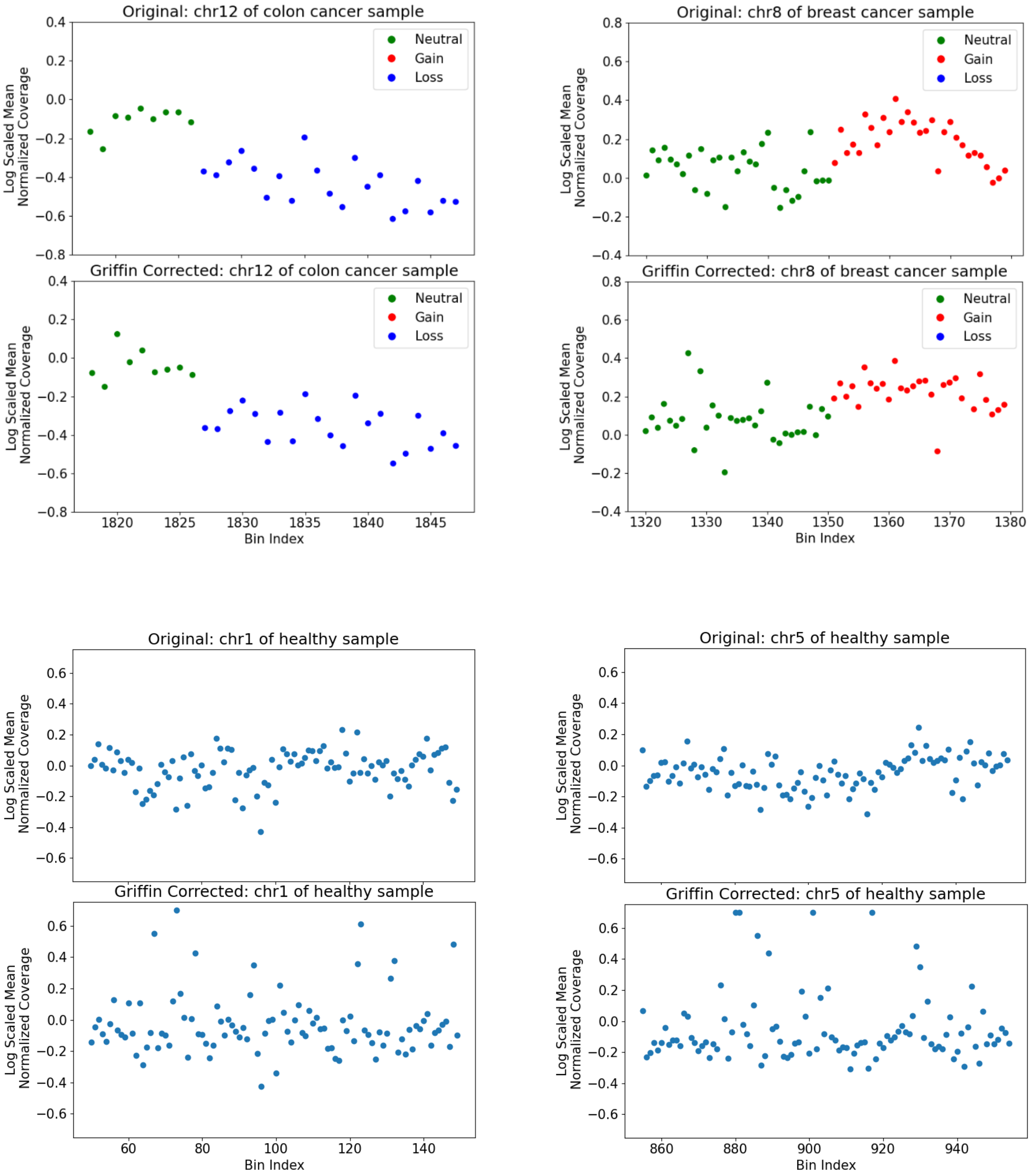


**Suppl Fig S13:** Griffin causes further disruption in the coverage profile for ultra low pass (~0.1X) WGS cancer and healthy samples

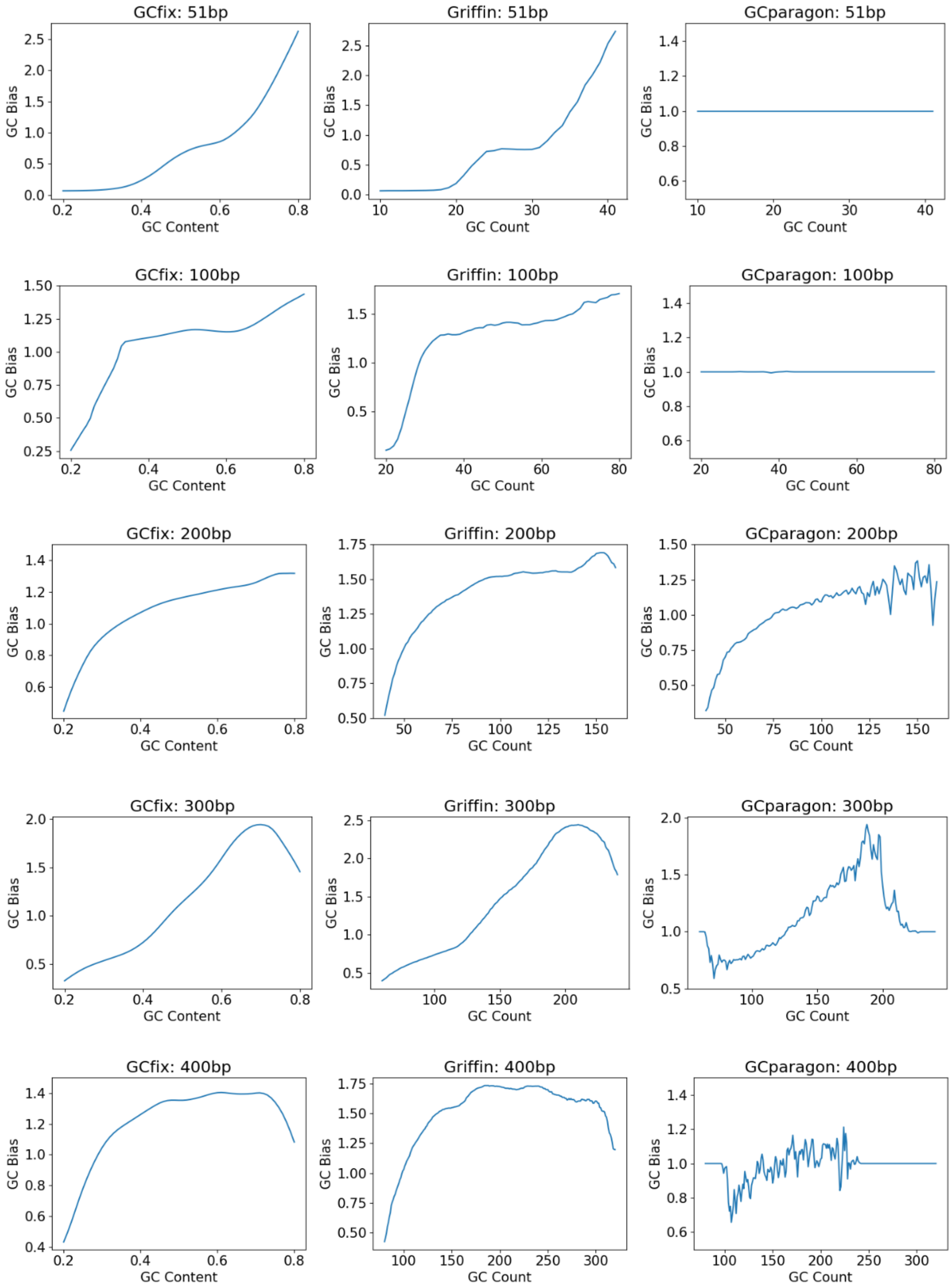


**Suppl Fig S14:** Comparing GC bias curves obtained from different GC correction methods for different fragment lengths of a healthy sample
