## Supplementary Data for "GCfix: A Fast and Accurate Fragment Length-Specific Method for Correcting GC Bias in Cell-Free DNA"

### **Supplementary Data 1:** Simulated Fetal Fraction Estimation Process

- The female pregnant BAM is made by mixing the full female BAM with X% male BAM; meaning that the mother is carrying X% male baby fetal fraction.
- All reads mapped to mother chrY should ideally come from male baby; meaning that $chrY_{frac}$ ($chrY_{read\_no}/chrY_{len}$) should come completely from male baby.
- We can calculate $chrN_{frac}$ for each autosome chrN of the mother using half of the reads mapped to that autosome (since we have a pair of autosomes): $chrN_{frac} = (chrN_{read\_no}/2)/chrN_{len}$
- $chrN_{frac} = \ chrY_{frac}$ only when 100% of mother chrN reads are coming from male baby denoting a fetal fraction of 100%
- We can estimate mother fetal fraction using mother autosome N using the formula: $chrY_{frac}/chrN_{frac}$
- Ideally, this estimated fetal fraction should be equal for all autosomes and they all should be similar as the ground truth fetal fraction.

Note that in order to obtain number of reads mapped to each chromosome, we only considered forward reads (to avoid double counting) with mapping quality of 60 or higher and excluded all supplementary, duplicate & unpaired reads from valid chromosome regions excluding blacklisted, unwanted and low mappability regions. Chromosome length is also calculated considering only the valid regions.

### **Supplementary Data 2:** Relative Coverage Calculation

#### Extraction of Sequencing Depth

Rsamtools package was used to extract per-read information from the sequencing data. Filtering was applied to retain only reads with fragment sizes between 51-400 bp, a mapping quality greater than 20, and reads aligned to the forward strand. Duplicate reads were also removed to prevent double counting.

Once the reads were filtered, the read depth was calculated. For original uncorrected read depth, a count of 1 was assigned per read. However, for GC-corrected read depth, the read count was adjusted based on the estimated GC-bias correction factor per read. This adjustment accounts for the inherent biases in sequencing related to GC content, providing a more accurate representation of the actual sequencing depth.

#### Plotting of Relative Coverage Profiles

The process of plotting relative coverage profiles shown in Suppl. Figure S11 involves several key steps. First, Nucleosome Depleted Region (NDR) site information is provided for the selected genes, along with the respective gene expression groupings. The coverage profiles are then extracted for each gene to proceed with the analysis.

Relative coverage is calculated by dividing the sequencing depth per base pair by the average sequencing depth of the flanking regions, specifically from +1000 to +2000 bp and -1000 to -2000 bp relative to the TSS. To prevent outliers from significantly skewing the results, the maximum relative coverage is capped at 2. Sites where the flanking region has an average sequencing depth of 0 are excluded from the comparison. For each base pair, the mean of the relative coverage is calculated for each sample. Subsequently, the relative coverage profiles are plotted at the cohort level, with the average relative coverage calculated per gene expression group across samples for visualization.

### **Supplementary Data 3:** Algorithmic Differences between GCfix, Griffin and GCparagon

- GCfix uses fixed 101 GC content % bins (0%, 1%, 2%, ..., 100%) for fragments of all lengths, whereas Griffin and GCparagon methods use raw GC counts. Possible raw GC counts vary widely depending on fragment length (length 51 can have counts of 0, 1, 2, ..., 51; length 350 can have counts of 0, 1, 2, ..., 350). As a result, these two methods cannot take advantage of parallelism and fast SIMD (Single Instruction, Multiple Data) instructions in different steps of GC correction.
- Griffin and GCparagon estimate GC correction factors at individual fragment length level. In our case, we merge frequencies of $\pm$2bp length fragments including the length of interest to estimate its GC correction factor; this gives us an advantage in lpWGS samples.
- Griffin and GCparagon use Gaussian/median smoothing on estimated GC bias values based on the GC bias values of fragments of neighboring lengths and similar GC contents. GCfix directly applies LOESS smoothing on fragment length specific GC bias values with a mild smoothing factor of 0.1.
- While existing methods use the full fragment length as the fragment specific GC context, we truncate 10 bp from both start and end of the fragment to avoid noisy behavior as shown in Suppl. Figure S1.
- Based on our analysis of deep and lpWGS samples coming from 4 different cohorts, we see that over 99% of the fragments come from 20-80% GC content context; hence GCfix provides GC correction factors for this range only making correction factors for GC content outside of this range to 0. Such measure is not taken in GCparagon and Griffin. GCparagon explicitly finds rare (fragment length, GC count) pairs to manually set their correction factors to 1. Griffin on the other hand does not perform any post processing of the estimated GC bias (inverse of bias is the correction factor). As a result, Griffin often produces NAN values and 0 values as their estimated GC bias for some combinations of GC count and fragment length.
- While GCparagon requires querying the reference genome for expected GC count frequencies every time one runs the method on a sample, GCfix requires doing this only once for a particular reference genome (Griffin also requires doing it only once).

- GCparagon uses a small subset of fragments and regions of deep WGS samples to estimate GC bias to keep the runtime under control; GCfix uses whole genome and all fragments to perform this estimation which is expected to be more accurate (Griffin also does the same thing).
- Griffin assumes that the GC bias curve for $\pm$10 bp length fragments are similar and so, it uses $\pm$10 bp length fragment GC bias values for smoothing GC bias values of the fragment of interest. This hypothesis is true only for fragments of size 100-200 bp inspected in the Griffin paper. But considering cfDNA fragment lengths of a wider range such as 51-400 bp, we found out that this similarity can be seen only up to $\pm$2 bp fragment length GC bias curves. As a result, as mentioned above, we merge frequencies of $\pm$2bp length fragments including the length of interest to estimate its GC correction factor.
- Griffin considers both forward and reverse reads for GC bias estimation; GCfix and GCparagon both utilize only forward reads.
- Griffin requires the users to create a suite of precomputed GC frequency file for the reference genome build the user wishes to use. The user requires a single expected GC frequency file per reference genome for GCfix. This file is already provided for hg19, GRCh37 and hg38. For a different reference genome, we have provided a single line command to generate this file.

### **Supplementary Data 4:** Obtaining Expected Fragment Count from Reference Genome

One key step of GCfix is to get the expected fragment counts for different GC contents for the different fragment lengths. Let us assume that we are deriving the expected counts for fragments of length 150. GC context for this fragment length is a window of size 130 excluding the first and the last 10 bp of the fragment. We take all possible positions of the valid genomic regions from the reference genome and assume that there is a fragment of size 150 in each of these positions. As a result, we would be looking at the GC context window of size 130 starting from 10 bp downstream of each position of the valid regions of the reference genome. For each position GC context window, we calculate the GC content of that window and increment the count of that particular GC content fragment number. After the entire process, we get the expected fragment count for different GC contents for fragment length 150. We can repeat the same process for fragments of different sizes using their specific GC context windows and get respective expected fragment counts.
